# A pooled CRISPR screen reveals genes critical for erythroblast enucleation

**DOI:** 10.64898/2026.04.06.716706

**Authors:** Marilou Tetard, Tianjian Lin, Nana A. Peterson, Rebekah C. Gullberg, Yann Le Guen, John G. Doench, Elizabeth S. Egan

## Abstract

Terminal erythroid differentiation involves dramatic cellular remodeling that culminates in the expulsion of the nucleus, a process known as enucleation. While enucleation is conserved across mammals and is crucial for the generation of fully functional erythrocytes, the mechanisms governing this process have remained largely unknown, in part because the absence of genetic material in mature, enucleated red blood cells hinders genetic experimentation. Here, we performed a pooled, forward-genetic CRISPR-Cas9 screen in enucleated red blood cells derived from primary human hematopoietic stem cells to identify genes required for enucleation. We found that Chloride Intracellular Channel 3 (CLIC3) and Vesicle-associated membrane protein 8 (VAMP8) are both necessary for terminal erythroid differentiation, yet likely act through different mechanisms. Knockdown of CLIC3 led to a delay in erythroblast differentiation, culminating in impaired enucleation. We found that the knockdown cells had increased p53 and p21 and exhibited cell cycle alterations, suggesting CLIC3 plays a crucial role in coordinating cell cycle progression during erythropoiesis. In comparison, VAMP8-depleted cells initially appear to undergo accelerated differentiation but then display a specific defect in enucleation. Transcriptional analysis of the VAMP8-knockdown cells suggested dysregulation of pathways for vesicle trafficking and actin binding, and imaging of late-stage erythroblasts revealed impaired nuclear polarization and disorganized actin. This work provides a new approach for functional genomics in enucleated cells and reveals novel factors important for terminal erythroid differentiation and enucleation.

**Key points:**

- A CROPseq-based CRISPR-Cas9 screen enables functional genomics in enucleated primary human red blood cells.
- Chloride Intracellular Channel 3 (CLIC3) and Vesicle Associated Membrane Protein 8 (VAMP8) were identified as critical for terminal erythroid differentiation and enucleation, likely acting through two distinct mechanisms.

## Introduction

Mammalian red blood cells (RBCs) are essential for oxygen transport and are continuously produced through erythropoiesis, a highly regulated process in which hematopoietic stem cells proliferate and differentiate down the erythroid lineage, yielding billions of new reticulocytes and erythrocytes every day^1^. Erythropoiesis unfolds in two distinct phases: an initial proliferation phase where hematopoietic progenitors commit to the erythroid lineage, followed by a terminal differentiation phase where erythroid precursor cells mature through a series of erythroblast developmental stages^2^. This pathway culminates in enucleation, a dramatic event in which the late-stage erythroblast expels its nucleus to become a flexible, enucleated reticulocyte^3^.

Enucleation is a complex process involving chromatin condensation and cell cycle exit, followed by organelle clearance, nuclear polarization and extrusion^4,5^. The force required for nuclear expulsion has been proposed to result from a measurable reorganization of the cytoskeleton, leading to a collection of F-actin at the rear of the cell, termed the enucleosome^6^. While these processes are known to involve various transcription factors, microRNAs, and cytoskeletal-associated proteins^5,7-9^, a comprehensive understanding of the molecular mechanism(s) driving enucleation is lacking.

Functional genetic approaches have proved fruitful for the discovery of genes important for erythropoiesis^10-13^, but less work has focused on enucleation, which is challenging to recapitulate in vitro. Available erythroid cell lines exhibit low rates of enucleation and may not model some critical biology of terminal erythropoiesis^14,15^; primary erythroblasts derived from hematopoietic stem/progenitor cells (HSPCs) enucleate more efficiently, opening the possibility of forward genetic screening for factors that govern this process^10,16,17^. However, the absence of nucleic acid in enucleated red cells presents challenges for large-scale forward-genetic screening, as tracking genetic perturbations in pooled screens typically relies on next-generation sequencing to quantify changes in the abundance of integrated sgRNA or shRNA sequences in different experimental conditions^18,19^. Here, we developed an approach for pooled CRISPR-Cas9 screening in enucleated human red cells derived from primary HSPCs to uncover new determinants of enucleation.

## Methods

A list of all reagents, antibodies, plasmids, sequences, and their sources can be found in supplemental Table 1. Additional methods can be found in “Supplemental Methods**”.**

### Ex-vivo erythropoiesis of primary human CD34^+^ HSPC

Primary human CD34^+^ HSPCs isolated from de-identified human bone marrow donors (Stem Cell Technologies, SCT) were cultured as previously described^17,20^, as detailed in Supplemental Methods. Briefly, cells were cultured in complete PIMDM (cPIMDM) supplemented with 3IU/ml Erythropoietin (Epo), 100 ng/ml Stem Cell Factor (SCF), 5 ng/ml IL-3, 10 µM hydrocortisone and 5% human plasma (Octapharma) at 37°C in 5% CO2 in air. IL-3 and hydrocortisone were removed on day 7, and the cells were maintained in fresh cPIMDM at a density below 5×10^5^ cells/ml. On day 11, SCF was removed and the cells were plated at 7.5 ×10^5^ - 1.0 ×10^6^ cells/ml in cPIMDM. On days 13–14, cells were plated on a murine stromal cell layer (MS-5) ^21^ at 1×10^6^ cells/ml.

### Pooled CRISPR screening

For each replicate, 1x10^6^ CD34^+^ primary human bone marrow HSPCs (SCT) were induced to undergo ex-vivo erythropoiesis as described above. On day 4, ∼1x10^7^ cells were transduced with the pooled lentivirus library. Details of the library design are described in supplemental Figure 2A. On day 5, every 1x10^7^ cells were nucleofected with 24.75µL of Cas9-NLS (40µM; Berkeley Macrolab) coupled with 8.25µL of a non-targeting ssODN (100uM) (Lonza Amaxa 4-D, program EO-100). Transduced cells were selected with puromycin from day 6 to day 11 and plated on a stromal cell layer from day 13 to day 16. Cells were harvested on day 8 (initial library representation) and day 17 (unsorted cells or FACS-sorted enucleated cells). RNA was extracted with a RNeasy Plus Mini Kit (Qiagen), with on-column DNAse, and eluted in 30µL. RNA quality was assessed using an Agilent 2100 Bioanalyzer. cDNA was produced from 24µL of RNA using SuperScript™ III (Invitrogen). sgRNA sequences were amplified by a 15-cycle PCR using primers Argon and Jeudi-Kermit (supplemental Table 1) and then sequenced at the Genetic Perturbation Platform of the Broad institute using Illumina sequencing.

### Measuring enucleation by flow cytometry

To quantify enucleated cells, differentiating cRBCs were incubated in Vybrant DyeCycle Ruby (1:10,000) in media at 37°C for 30 min in the dark. Flow cytometry was performed on a MACSQuant (Miltenyi Biotec).

### CRISPR screen analysis

Enrichment at the sgRNA and gene levels was determined using developed by the Genetic Perturbation Platform at the Broad institute (Apron version 1.1.0). After normalization, a *z*-score was calculated for each sgRNA from log_2_-fold changes between the unsorted and enucleated populations using the standard deviation and mean of non-targeting guides^22^. To obtain z-scores at the gene level, for each condition z-scores of all constructs for a given gene were summed and divide by the square root of the standard deviation of number of constructs for this target (Stouffer’s method). *P*-values were calculated directly from the gene z-scores using the standard distribution on the average log-fold changes. The false discovery rates were calculated from *P*-values using the Benjamini-Hochberg procedure.

### Genetic modification of primary human HSPCs with individual shRNAs

CD34+ human HSPCs from bone marrow were cultured as described in supplemental Methods. On day 3 or 5 of differentiation 1x10^6^ cells were transduced with lentivirus with 4µg/ml polybrene via spinoculation at 1000xg for 2h. Transduced cells were selected with puromycin.

### Quantification of mitochondria by flow cytometry

To quantify mitochondria, day 13 cRBCs were incubated with 100nM MitotrackerGreen (ThermoFisher) for 15 minutes at 37°C in the dark. After two washes in PBS, cells were staged using anti-Band 3-PE anti CD49d-APC as described in supplemental Methods and finally incubated with Hoechst (1:5000) for 10 minutes. Flow cytometry was performed on a MACSQuant (Miltenyi Biotec).

### Cell Cycle assays

Cell cycle assays were performed using the Click-iT™ Plus EdU Alexa Fluor™ 647 Flow Cytometry Assay Kit (Thermo) according to the manufacturer’s instructions. Briefly, 2x10^5^ day 7 cRBCs were incubated with 10 µM EdU for 1 hour at 37°C and then treated following the Click-iT kit protocol. After washing, cells were incubated in DAPI (1 µg/mL) for 15 minutes protected from light. Flow cytometry was performed on a MACSQuant (Miltenyi Biotec).

### Fluorescence microscopy

Immunofluorescence Assays (IFAs) for localization of CLIC-3 and VAMP-8 were performed using standard methods, as described in “supplemental Methods”. Nuclear polarization was quantified for each genetic background by blindly counting all cells in 9 fields of view (>700 nucleated cells per condition). Actin foci were quantified by blindly counting all cells across 9 fields of view (>800 cells per condition). IFA images were acquired using Zeiss LSM 980 with Airyscan 2.

### Data Analysis

All flow cytometry data were analyzed using FlowJo (v.10.8.1). Confocal images were analyzed with Zeiss Zen 3.11, and FIJI (v.2.9.0). Western Blots were analyzed using the Odyssey Fc imager (LI-COR) and quantified using Image Studio software (LICORbio). Bulk RNA sequencing was performed on a NovaSeq X Plus Series (PE150). Differential gene expression analysis was performed using DESeq2. Gene-wise counts were modeled using a negative binomial framework, and *P*-values were adjusted for multiple testing using the the Benjamini-Hochberg method to control the false discovery rate (FDR). Pathway analysis was performed using ShinyGO.0.85.1^23^, using the Hallmark MSigDB database and the GO:cellular component database. Statistical analyses were performed with GraphPad Prism version 10.

## Results

### A method for pooled CRISPR screening in enucleated red blood cells

To uncover genetic determinants of enucleation, we established a pooled CRISPR screening strategy in cultured red blood cells (cRBCs) derived ex-vivo from primary human CD34^+^ HSPCs isolated from bone marrow. Our approach for ex-vivo erythropoiesis routinely yields ∼10,000-fold expansion with >90% enucleation after ∼18-20 days, producing large numbers of enucleated cRBCs (Figure 1A-C).

**Figure 1.**
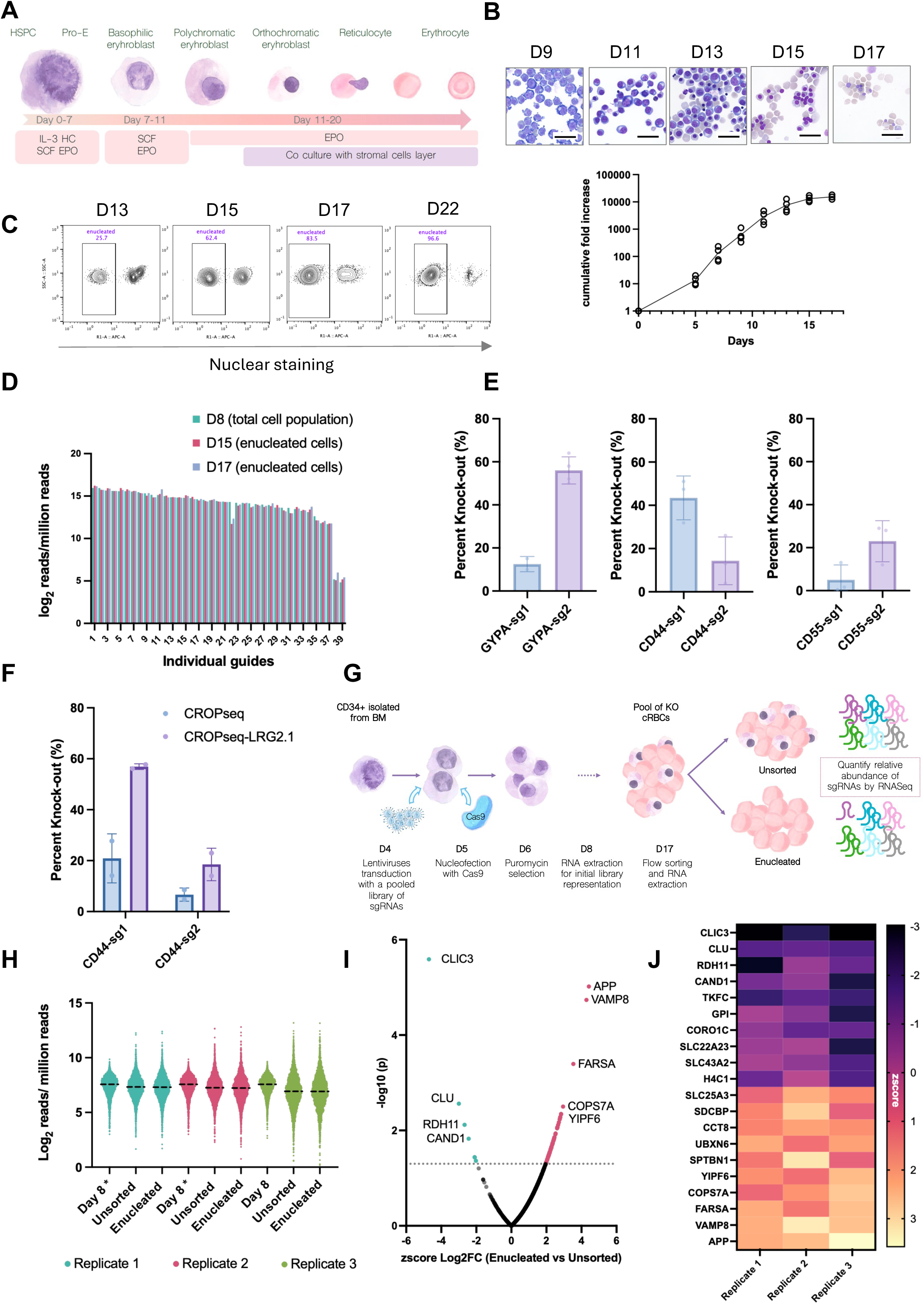
Development of a pooled CRISPR screen for enucleation factors in human red blood cells. (A) Schematic representation of ex-vivo erythropoiesis approach to generate cRBCs from primary human CD34^+^ HSPCs. (B) Representative cytospin images of differentiating erythroid precursor cells over a time-course of ex-vivo erythropoiesis; scale bar, 30 μm. Growth curves of primary human bone marrow CD34^+^ HSPCs during ex-vivo erythropoiesis. The line shows the mean cumulative fold increase in growth, *n*=4 biological replicates. (C) Representative flow cytometry plots showing enucleation rate of cRBCs during terminal erythroid differentiation, as detected by flow cytometry using a cell-permeable DNA stain. (D) Bar plot showing the relative abundance of 39 CROPseq-encoded sgRNAs in differentiating cRBCs as quantified by RNA-seq. For day 15 and 17 cells, RNA was prepared from enucleated cRBCs sorted by FACS. (E) Efficiency of gene knockout in differentiating cRBCs after transduction of CROPseq lentivirus expressing indicated sgRNAs followed by Cas9-NLS nucleofection (SLICE). Percent knockout was measured relative to mock cRBCs via flow cytometry for surface protein expression. Data points represent the mean ± SD of at least 2 biological replicates. (F) Efficiency of gene knockout in cRBCs via SLICE using the original CROPseq vector or a modified vector containing the scaffold from LRG2.1(CROPseq-LRG2.1). Percent knockout was measured relative to Mock cRBCs via flow cytometry. Bars show the mean ± SD; *n*=2. (G) Schematic of strategy for pooled CRISPR-Cas9 screen in primary human HSPCs. CD34^+^ HSPCs isolated from bone marrow were induced to undergo ex-vivo erythropoiesis, transduced with a custom CROPseq lentivirus library on day 4 and nucleofected with rCas9-NLS on day 5. On day 17, enucleated cRBCs were sorted by FACS. sgRNA sequences were quantified by RNA-seq on day 8 (baseline) and in two populations on day 17 (unsorted cells and FACS-sorted enucleated cells). (H) Library distribution in cRBCs at indicated timepoints. Each data point represents a unique sgRNA sequence embedded within the mRNA, as detected by RNA-seq. Dotted line represents the median sgRNA relative abundance for each population; *\** identical data set. (I) Volcano plot of the combined 3 replicates showing gene-level representation in enucleated versus unsorted cells on day 17. The x-axis represents the *z*-score of the log_2_ fold change (log2FC), and the y-axis represents the -log10(*P* -value). Genes with sgRNAs significantly decreased in relative abundance in enucleated cells are in blue, and genes with sgRNAs increased in enucleated cRBCs are in pink. (J) Heatmap showing the *z*-score for the top 10 enriched and depleted hits across three biological replicates.

A potential challenge for CRISPR-based pooled screening for determinants of enucleation is that single guide RNAs (sgRNAs) may not be exported out of the nucleus nor detectable after enucleation. Indeed, we found that sgRNAs expressed from a U6 lentivirus promoter were not detectable in enucleated cRBCs even though they were detected in late-stage nucleated erythroblasts (supplemental Figure 1A). To overcome this, we adapted the CROPseq vector, in which the sgRNA cassette is inserted into the 3’ LTR, leading to its incorporation within a polyadenylated mRNA transcript that may be exported to the cytoplasm^24^ (supplemental Figure 1B). As a proof of principle, we transduced erythroid progenitors generated from primary CD34^+^ HSPCs with CROPseq lentivirus expressing an sgRNA targeting glycophorin A (GYPA). After differentiation and sorting for enucleated cRBCs, we observed that the sgRNA was detectable by RT-PCR post-enucleation, indicating that mRNA harboring the embedded sgRNA sequence is exported to and retained in the cytoplasm (supplemental Figure 1C).

To validate this approach for a pooled screening format, we introduced a pilot CROPseq library of 39 sgRNAs^25^ into HSPCs and induced them to differentiate down the erythroid lineage. RNA-seq analysis of the population at day 8, and of the enucleated populations at days 15 and 17, revealed that the relative abundance of each sgRNA remained stable as the cells matured and enucleated (Figure 1D; supplemental Figure 1D). Together, these results establish the feasibility of quantifying CROPseq-encoded sgRNAs in enucleated cRBCs.

To optimize editing efficiency for a pooled screen, we adapted a strategy known as SLICE (sgRNA lentiviral infection with Cas9 protein electroporation), in which transduction of a lentivirus CROPseq library is followed by nucleofection of rCas9-NLS^26^. To test this approach in primary erythroid cells, we transduced day 4 cells with individual CROPseq lentiviruses targeting the RBC surface proteins CD44, CD55, or GYPA and introduced rCas9 protein by nucleofection the following day. This approach led to varied knockout efficiencies depending on the sgRNA (Figure 1E). However, replacing the CROPseq sgRNA scaffold with that of vector LRG2.1, which is optimized for CRISPR-knockout^27^, substantially improved the editing efficiency (Figure 1F). Thus, SLICE with an optimized CROPseq construct (CROPseq-LRG2.1) enables efficient and traceable gene editing in enucleating cRBCs.

### Pooled CRISPR-Cas9 screen reveals genes necessary for enucleation

To screen for determinants of erythroblast enucleation, we designed a custom CROPseq-LRG2.1 sgRNA library targeting the RBC proteome. We curated a list of 681 genes encoding membrane, cytoskeletal, and signaling proteins from published RBC proteomic datasets^28-30^, excluding known essential genes and those with previously described roles in erythropoiesis (supplemental Figure 2A). The library consisted of 5,007 sgRNAs (7 per gene plus 240 non-targeting controls). The screen was performed in three biological replicates, using bone marrow CD34^+^ HSPCs from two de-identified donors (supplemental Table 2). HSPCs were transduced with the sgRNA library on day 4, nucleofected with rCas9-NLS the following day, and induced to differentiate down the erythroid lineage (Figure 1G). Cells were harvested for initial library representation on day 8. Cells were again harvested on day 17, and enucleated cells were isolated by FACS. mRNA was isolated from all three populations (day 8, day 17 unsorted, and day 17 enucleated) and the relative abundance of each sgRNA sequence embedded in total mRNA was quantified by RNA-sequencing (supplemental Table 2).

Analysis of sgRNA abundance revealed a broad and unimodal distribution of the library at each of the three time points, confirming that the library complexity was preserved throughout the experiment (Figure 1H). To identify candidate genes for enucleation, we quantified the relative abundance of guides in each population normalized to the non-targeting guides using a custom algorithm previously described^22^ (Figure 1I; supplemental Figure 2B; supplemental Table 3-4). Among the highly ranked genes, Chloride Intracellular Channel 3 (CLIC3) arose as a top candidate due to a marked reduction in sgRNA abundance in enucleated versus unsorted cells (Figure 1I-J; supplemental Figure 2C-F). Some CLIC3 sgRNAs were also relatively depleted on enucleated cells on day 17 as compared to the baseline population on day 8, suggesting a possible broader role for this gene in terminal erythroid differentiation (supplemental Figure 2C, E). Notably, another highly ranked candidate, retinol dehydrogenase (RDH11), has previously been shown to accelerate cell proliferation in an erythroleukemia cell line, consistent with a role in erythropoiesis^31^ (Figure 1I-J).

The screen also identified sgRNAs that were relatively overrepresented in the enucleated population, suggesting that the genes targeted by these sgRNAs may negatively regulate enucleation (Figure 1I-J, supplemental Figure 2G-H). One of the top candidates, Vesicle associated membrane protein 8 (VAMP8), is an integral membrane protein in the SNARE family of proteins, which are essential for fusion of cellular membranes. To our knowledge, neither CLIC3 nor VAMP8 have been previously implicated in terminal erythroid differentiation or enucleation.

### CLIC3 is required for erythroid enucleation

CLIC3 is associated with a variety of malignancies, with proposed roles ranging from acting as a chloride channel, a secreted oxidoreductase, an epigenetic regulator of cell cycle genes, and as an enhancer of integrin recycling to the plasma membrane^32-35^. To investigate a potential role in erythropoiesis, we first assessed CLIC3 expression in a time course and found that it is expressed throughout terminal erythroid differentiation, with protein levels remaining relatively stable while mRNA levels increased, suggesting a potential role in terminal differentiation (Figure 2A-B). Immunofluorescence microscopy revealed that CLIC3 localized to both the cytoplasm and nucleus in early erythroblasts, becoming predominantly cytoplasmic in later stages (Figure 2C).

**Figure 2.**
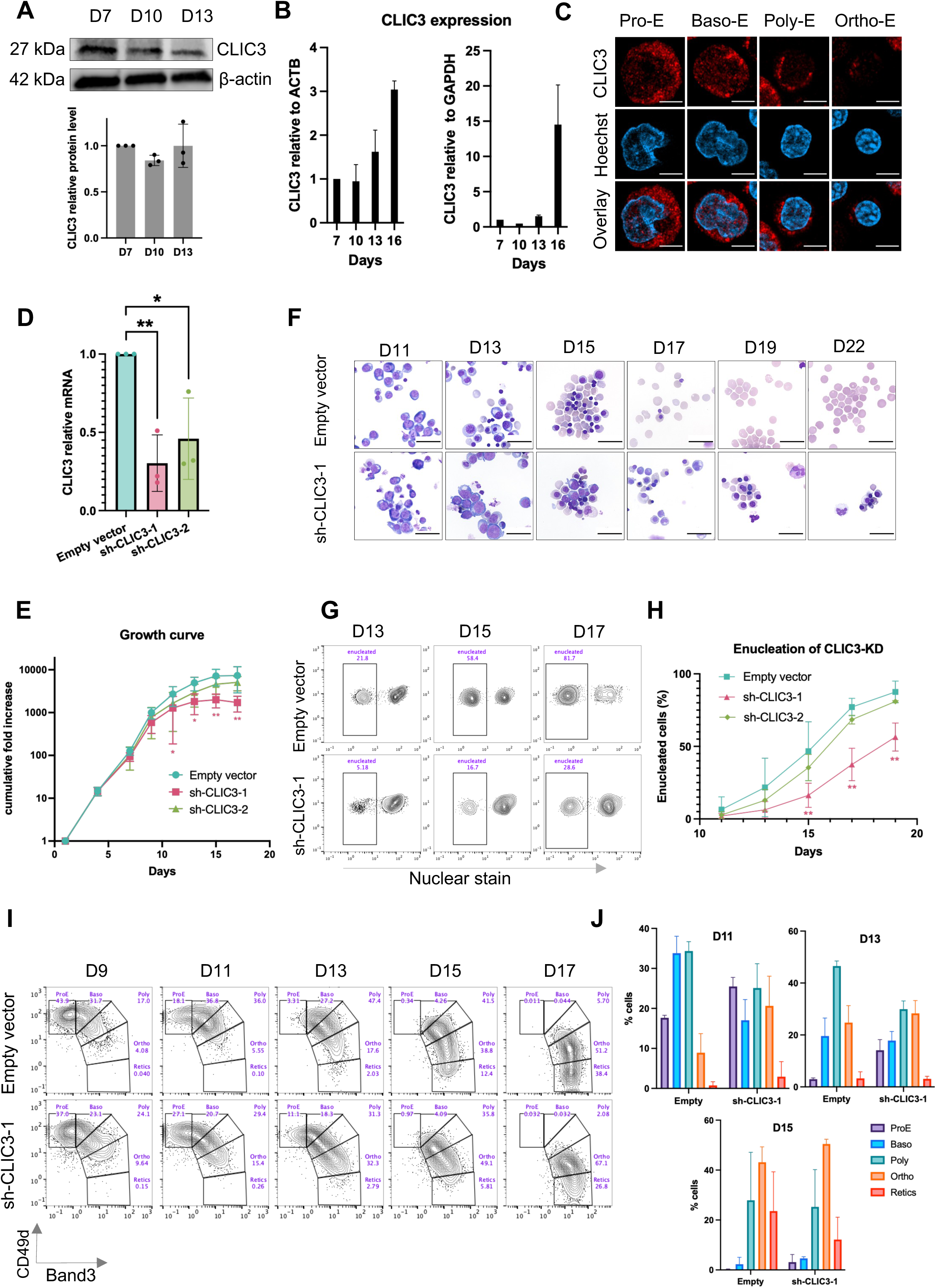
CLIC3 is required for terminal erythroid differentiation and enucleation. (A) Western blot to quantify CLIC3 in erythroblast lysates at day 7, day 10 and day 13 of differentiation. β-actin was used for normalization. Barplot shows the quantification of CLIC3 relative protein level normalized on β-actin of *n*=3 biological replicates. (B) Bar plots of CLIC3 mRNA levels at different days as measured by qRT-PCR. Data are either normalized to *ACTB* (left) or to *GAPDH* (right) and expressed relative to the mRNA level on Day 7. Values are mean ± SD of n=3 biological replicates. (C) Immunofluorescence assay for the localization of CLIC3 (red) in differentiating erythroblasts, with nuclei stained with Hoechst (blue). Scale bar, 5 μm. (D) Bar plot of CLIC3 mRNA levels in day 11 CLIC3-KD (sh-CLIC3-1 or sh-CLIC3-2) and isogenic WT erythroblasts (Empty vector) as measured by qRT-PCR. Data were normalized to GAPDH and expressed relative to WT. Bars show mean ± SD, *n*=3. *P* -values were calculated using an unpaired t test (\**p* <0.05; \*\**p* < 0.01). (E) Growth curves of CD34^+^ HSPCs with knockdown of CLIC3 (sh-CLIC3-1, sh-CLIC3-2) or isogenic WT (Empty vector) during erythroid differentiation. Data points represent the mean ± SD of 4 biological replicates. *P*-values were calculated using a Mann-Whitney test (\**p* <0.05; \*\**p* < 0.01). (F) Representative images of May-Grünwald- and Giemsa-stained cytospins from isogenic WT (Empty vector) or CLIC3-KD (sh-CLIC3-1) primary erythroid cells during a time course of terminal differentiation. Scale bar, 30 μm. (G) Representative experiment showing quantification of enucleation in isogenic WT (Empty vector) or CLIC3-KD (sh-CLIC3-1) cRBCs by flow cytometry. (H) Time course of erythroid enucleation in WT or CLIC3-KD cRBCs as measured by flow cytometry. Values represent the mean ± SD of *n*=3 biological replicates. *P*-values were calculated using a Mann-Whitney test (*p <0.05; **p < 0.01). (**I**) Flow cytometric analysis of erythroblast maturation via surface staining for α4-integrin and Band3 in cells pre-gated for GYPA expression. α4-integrin decreases during erythroid maturation whereas Band3 increases, as indicated by gating for erythroblast stages. (J) Quantitative analysis of the different stages based on the gating strategy showed on Fig 2-I during terminal erythropoiesis in cells transfected with an empty vector and cells transfected with a sh-CLIC3-1. Values represent the mean ± SD of *n*=3 biological replicates.

To validate a role for CLIC3 in enucleation, we used two independent shRNAs to deplete CLIC3 via RNAi, leading to a 50-70% reduction in mRNA (Figure 2D). Upon induction of ex-vivo erythropoiesis, CLIC3-knockdown (CLIC3-KD) cells displayed a mild yet reproducible proliferation defect, yielding up to ∼4-fold fewer cells by day 17 as compared to isogenic control cRBCs (Figure 2E). Morphological analysis revealed a striking enucleation phenotype by day 17, as the WT cRBCs were predominantly enucleated, but the isogenic CLIC3-KD cRBCs were predominantly nucleated late-stage erythroblasts (Figure 2F). The impaired enucleation in the CLIC3-KD cells was confirmed by quantitative flow cytometry, revealing ∼80% enucleation in WT cRBCs vs ∼30% in CLIC3-KD on day 17 (Figure 2 G-H).

Since our CRISPR screening data suggested that some CLIC3-targeting sgRNAs were decreasing in abundance in day 17 relative to day 8 cRBCs, we sought to determine whether earlier stages of erythroid differentiation might also be impacted by CLIC3 knock-down. Analysis of cytospin slides revealed morphological differences in the populations as early as day 11, with an apparent delay in the development of the CLIC3-KD cells (Figure 2F). Flow cytometry analysis of surface markers of erythroid differentiation confirmed this observation, with higher numbers of proerythroblasts in the CLIC3-KD cells on day 11 and 13, and a slower trajectory of differentiation as compared to isogenic WT cRBCs through day 15 (Figure 2I-J). Collectively, these results show that CLIC3 is required for efficient erythroid proliferation, terminal differentiation, and enucleation.

### CLIC3 knockdown leads to perturbation of the cell cycle

To gain insight into the molecular mechanism underlying the CLIC3-KD phenotype, we performed RNA-sequencing (RNA-seq) on isogenic WT and CLIC3-KD erythroblasts on day 11. This revealed hundreds of differentially expressed genes, including downregulation of genes encoding hemoglobin subunits and late-stage erythroid markers (e.g. *HBM, HBB, SLC4A1*), consistent with the observed developmental delay (Figure 3A-B, supplemental Table 5). Pathway analysis showed that *CLIC3* knockdown was associated with dysregulation of multiple pathways, including those implicated in cell cycle regulation, inflammatory responses, and epithelial mesenchymal transition, a hallmark of differentiation (Figure 3C). One notably dysregulated pathway was the p53 transcriptional gene network, which has a well-described role in terminal erythroid differentiation^36,37^ (Figure 3C; supplemental Figure 3A-C). Among the most significantly upregulated genes in this pathway was *CDKN1A/P21*, which is known to be upregulated in ineffective erythropoiesis^38,39^. We confirmed this result by both qRT-PCR and Western blot, demonstrating significantly increased p21 mRNA and protein levels in CLIC3-KD cells (Figure 3D-E). We also observed an increase in p53 protein levels, although no changes in RNA were observed, suggesting CLIC3-KD may be associated with increased p53 activation (Figure 3E; supplemental Figure 3C). Notably, no change was observed in protein abundance nor in RNA transcripts for the master erythroid regulator GATA-1, suggesting that the p21 induction observed in CLIC3-KD cells occurs via a GATA-1-independent mechanism (Figure 3E).

**Figure 3.**
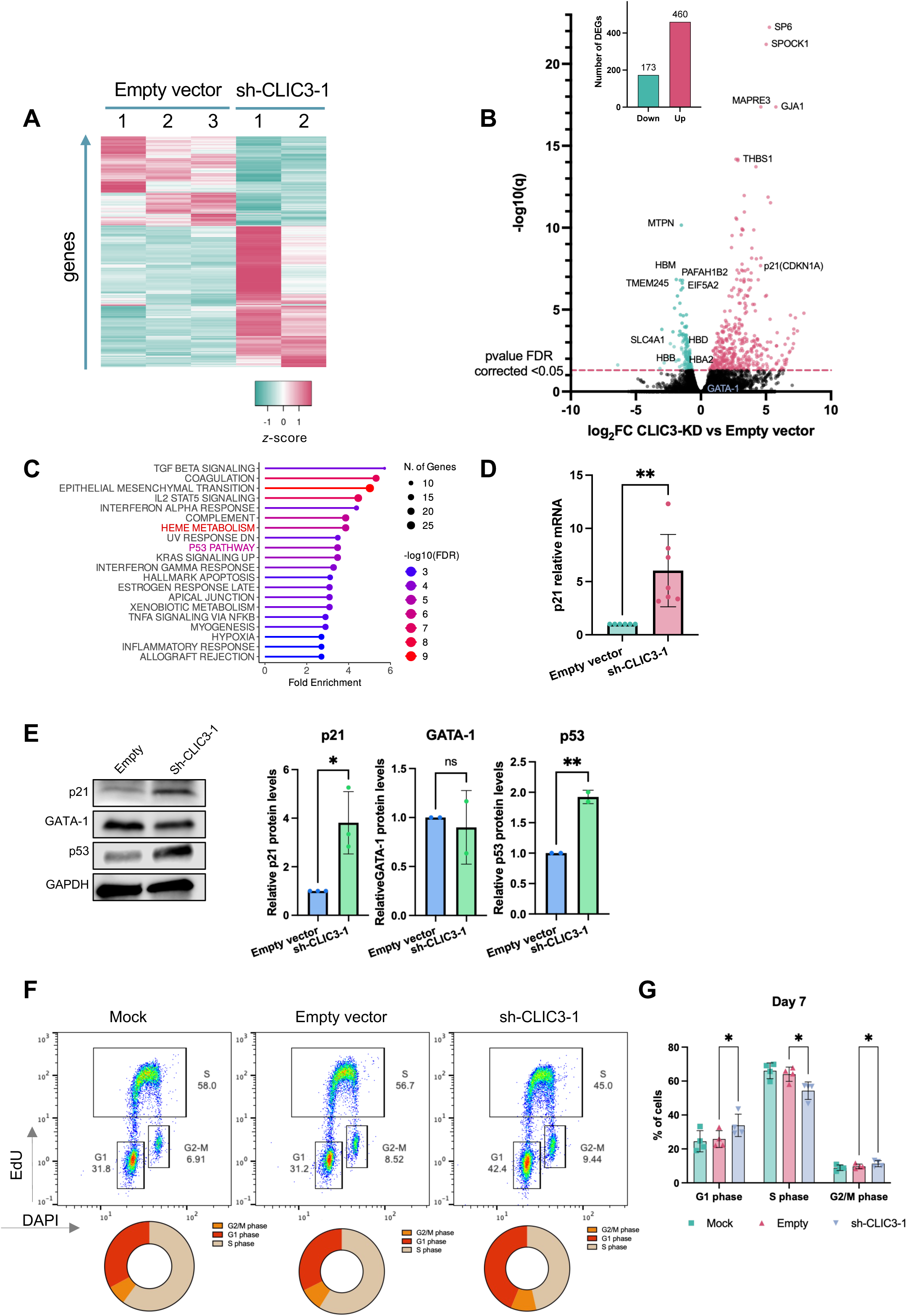
CLIC3 knockdown induces cell cycle dysregulation in erythroblasts. (A) Heatmap depicting differentially expressed genes in WT (Empty vector) versus CLIC3-KD (sh-CLIC3-1) day 11 erythroblasts. The color scale indicates *z*-score of gene expression. (B) Volcano plot showing differentially expressed genes between WT and CLIC3-KD erythroblasts on day 11. 460 genes are up regulated in CLIC3-KD, and 173 genes are down regulated (*P*-value FDR corrected <0.05). *n*= 3 for WT and *n*=2 for CLIC3-KD. Dotted lines indicate a threshold of FDR < 0.05. (C) Pathway analysis showing enriched gene sets from the MSigDB Hallmark database for differentially expressed genes between WT and CLIC3-KD erythroblasts on day 11. (D) Bar plot of *p21* mRNA levels in CLIC3-KD and isogenic WT, day 11 erythroblasts, as measured by qRT-PCR. Values are mean ± SD, *n*=7 biological replicates (** *P*-value <0.01). Data were normalized to *GAPDH* and expressed relative to WT. (E) Representative western blot showing p21, p53 and GATA-1 protein levels in isogenic WT versus CLIC3-KD erythroblasts on day 11 of differentiation. Bar plot represents protein level relative to WT, normalized to GAPDH; *n*=2-3 biological replicates *P*-values were calculated using an unpaired t test (* *P*-value <0.05; ** *P* -value <0.01). (F) Representative experiment showing cell cycle status of WT versus CLIC3-KD erythroblasts on day 7 of differentiation, as measured flow cytometry for EdU incorporation and DNA abundance (Dapi). (G) Quantitative analysis of the cell cycle phases for isogenic WT versus CLIC3-KD erythroblast populations on day 7 of differentiation, based on the gating strategy showed in Figure 3F. Bars indicate mean ± SD, *n*=4 biological replicates. *P*-values were calculated using paired t-test (* *P*-value <0.05)

Increased p53 and p21 have been implicated in the pathology of congenital anemia, where they are associated with cell cycle arrest in erythroid progenitor cells^39^. To functionally examine the cell cycle in CLIC3-KD, we performed an EdU incorporation assay on day 7, when the erythroblasts are still highly proliferative. Quantitative analysis revealed that CLIC3-KD resulted in a reduced percentage of cells in S-phase (45% vs 57%), while the percent of cells in G1 was significantly increased (42% vs 31%) (Figure 3F-G). Taken together, these results suggest that CLIC3 depletion triggers p53/p21-dependent cell cycle dysregulation, leading to delayed differentiation and impaired enucleation. Alternatively, CLIC3 may play multiple, distinct roles during the multi-step process of erythroid differentiation, impacting the cell cycle and enucleation separately.

### VAMP8 is required for enucleation

VAMP8 arose as an additional top candidate from our screen, with the prediction (based on an enrichment of *VAMP8*-targeting sgRNAs in the enucleated cRBCs) that knockdown would enhance enucleation (Figure 1I-J; supplemental Figure 2G-H). VAMP8 is a member of the SNARE family of proteins, which mediate the fusion of vesicles with target membranes and play key roles in diverse biological processes^40,41^. In most cell types, VAMP8 is localized to the lysosomal membrane and has been shown to regulate lysosome–autophagosome fusion^42^.

As VAMP8 has not been characterized in erythroid cells, we first determined its expression and localization in terminally differentiating human erythroblasts. Western blot analysis showed stable VAMP8 protein expression during differentiation, relative to GAPDH (Figure 4A). Immunofluorescence assays revealed a primarily cytoplasmic distribution with a punctate appearance for VAMP8 starting from the pro-erythroblast stage, with partial colocalization with the lysosomal marker LAMP2 (Figure 4B).

**Figure 4.**
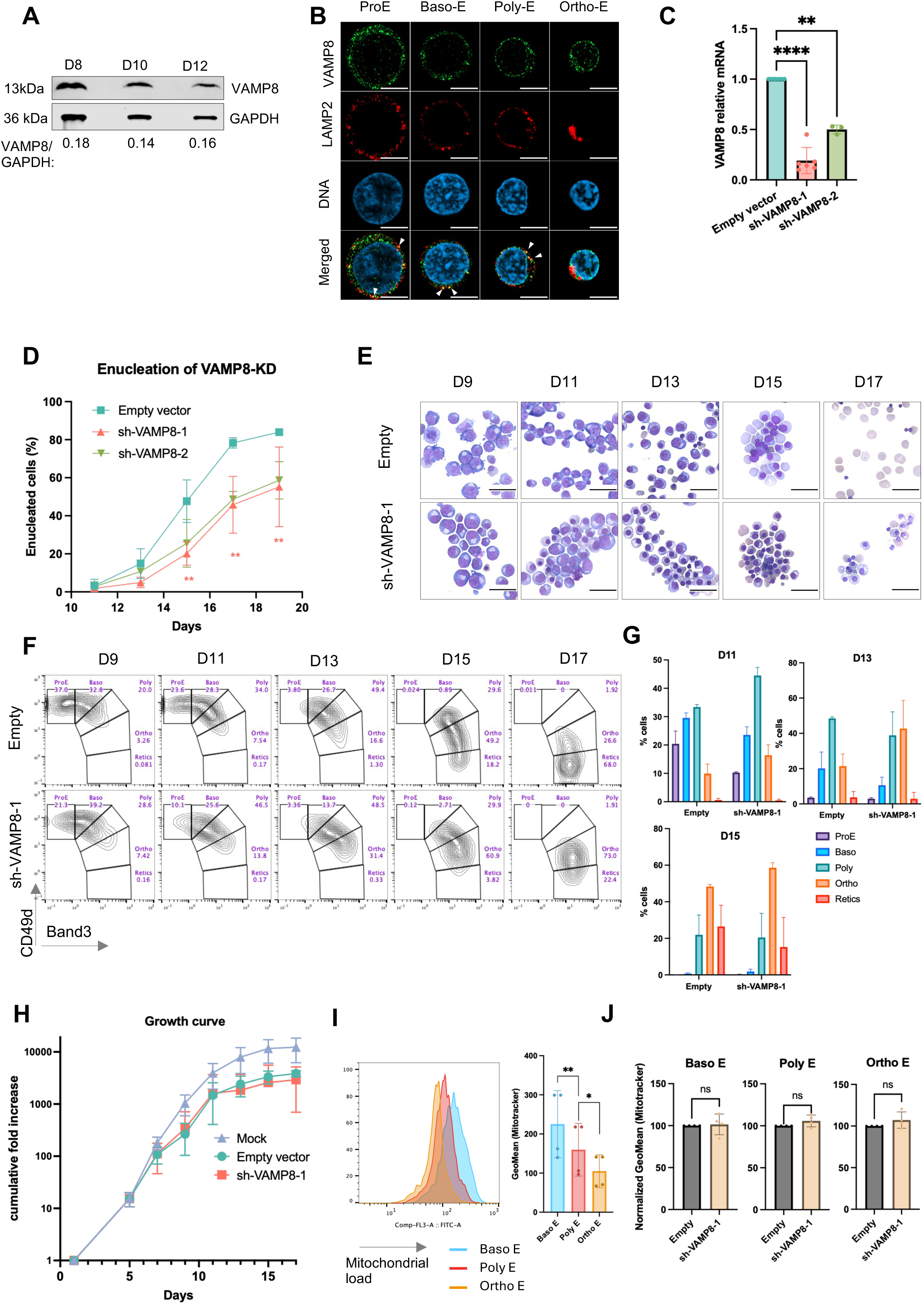
VAMP8 is required for erythroblast enucleation. (A) Western blot to quantify VAMP8 in erythroblast lysates at day 8, day 10 and day 12 of differentiation. GAPDH was used for normalization. Ratio between VAMP8 and GAPDH signal is indicated below. (B) Immunofluorescence assay showing the localization of VAMP8 (green), LAMP2 (red) and nuclei (blue) in differentiating erythroblasts. Arrows indicate regions of colocalization. Scale bar, 5μm. (C) Bar plot of *VAMP8* mRNA levels in isogenic WT (Empty vector) or VAMP8-KD (sh-VAMP8-1 or sh-VAMP8-2) day 11 erythroblasts as measured by qRT-PCR. Data were normalized to *GAPDH* and expressed relative to WT. Values are mean ± SD at least three replicates. *P*-value were calculated using a t-test (** *P*-value <0.01; **** *P*-value<0.0001). (D) Time course of enucleation of isogenic WT or VAMP8-KD cRBCs, as measured by flow cytometry. Values represent the mean ± SD of *n*=3 biological replicates. *P*-values were calculated using a Mann Whitney test (** *P*-value <0.01). (E) Representative cytospin images of a time course of erythroid differentiation of WT and isogenic VAMP8-KD erythroblasts. Scale bar, 30 μm. (F) Representative flow cytometry experiment to analyze differentiation of isogenic WT of VAMP8-KD erythroblasts via surface staining for α4-integrin and Band3 after pre-gating for GYPA expression. α4-integrin decreases during erythroid maturation whereas Band3 increases, as indicated by gating for erythroblast stages. (G) Quantitative analysis of the different stages based on the gating strategy showed on Figure 4F. Values represent the mean ± SD of *n*=3 biological replicates. (H) Growth curves of CD34^+^ HSPCs with knockdown of VAMP8 (sh-VAMP8-1) or isogenic WT (Empty vector) during erythroid differentiation. Data points represent the mean ± SD of 3 biological replicates. *P*-values were calculated using a Mann-Whitney test, *P*=n.s. (I) Representative histograms of mitochondrial abundance in WT basophilic (Baso E), polychromatic (BasoE) and orthochromatic (Ortho E) erythroblasts as measured by Mitotracker staining and analyzed by flow cytometry. Gating for erythroblast populations was based on a4-integrin/Band3 profiles as shown in Figure 4F, with inclusion of nuclear stain to gate out enucleated cells. Barplot shows the GeoMean of Mitotracker in Baso E, Poly E and Ortho E. Values are mean ± SD of *n*=4 biological replicates. *P*-value were calculated using t-tests (** *P* -value <0.01; * *P* -value<0.05). (J) Barplot showing the quantification of mitochondrial abundance isogenic WT or VAMP8-KD basophilic (Baso E), polychromatic (Poly E) and orthochromatic (Ortho E) erythroblasts, as measured by Mitotracker staining and flow cytometry. Values were normalized to the mitochondrial abundance in the WT condition, and bars show the mean ± SD; *P*-values were calculated using a paired t-test *n*=4 biological replicates.

To validate the screen result and assess a potential role for VAMP8 in erythroid enucleation, we generated VAMP8-knockdown (VAMP8-KD) in cRBCs using two independent shRNAs. On day 11, we observed *VAMP8* knock-down efficiencies of ∼80% and ∼50% respectively (Figure 4C). Next, to determine the impact of VAMP8 on enucleation, we induced the erythroid precursor cells to undergo terminal differentiation and quantified the percent of enucleated cells by flow cytometry. Depletion of VAMP8 revealed a marked enucleation defect, with a ∼40% reduction in enucleation compared to control cells by day 19 (Figure 4D). This phenotype was confirmed by microscopy, showing an accumulation and persistence of late-stage, nucleated orthochromatic erythroblasts in the VAMP8-KD population (Figure 4E). Notably, examination of the cellular morphology at earlier timepoints suggested that the VAMP8-KD cells may initially have an accelerated rate of maturation. This was confirmed via flow cytometry of differentiation markers, which showed that by day 11, VAMP8-KD cRBCs had a ∼2-fold enrichment in orthochromatic erythroblasts compared to controls, a trend that was maintained at day 13 (Figure 4F-G). Unlike for CLIC3, VAMP8-KD had no apparent effect on cell proliferation (Figure 4H). These findings suggest a necessary and specific role for VAMP8 in erythroid enucleation.

### VAMP8-KD shows no detectable defect in mitochondrial clearance

The steps of terminal erythroid differentiation involve cell cycle arrest, chromatin condensation, polarization of the nucleus, and progressive organelle clearance. The VAMP8-KD phenotype of accelerated differentiation followed by impaired enucleation is reminiscent of that described for cRBCs with knockdown of the outer mitochondrial membrane protein voltage-dependent anion channel-1 (VDAC1)^43^. During erythroid differentiation, VDAC1-KD cRBCs fail to clear mitochondria efficiently, leading to accumulation of abnormal mitochondria and cell death^43^. As VAMP8 has been implicated in autophagy in other cells, we questioned whether the enucleation phenotype may be linked to a defect in mitochondrial clearance (mitophagy). As reported previously^43^, during the transition from basophilic to polychromatic to orthochromatic erythroblasts, a decrease in mitochondrial staining can be observed by flow cytometry, reflecting progressive clearance of mitochondria (Figure 4I; supplemental Figure 4A). However, there were no significant differences in the abundance of mitochondria in VAMP8-KD versus isogenic WT cRBCs at any of the stages of terminal erythroid differentiation, suggesting that unlike for VDAC1, VAMP8 is not required for mitochondrial clearance prior to enucleation (Figure 4J, supplemental Figure 4A-B).

### VAMP8 knock-down impacts actin reorganization during enucleation

To investigate the molecular basis for the enucleation block in VAMP8-depleted cRBCs, we performed RNA-seq on isogenic WT and VAMP8-KD erythroblasts at day 11. Differential expression analysis identified 400 differentially expressed genes (*P*-value <0.05), including downregulation of VAMP8 (Figure 5A-B; supplemental Table 6). However, only a modest number of differentially-expressed genes reached FDR significance, likely reflecting inter-replicate variability in these primary cells. Notably, the gene with the most significance differential expression was the actin binding protein ABLIM1. When considering all differentially-expressed genes, gene Ontology (GO) analysis revealed an enrichment for terms related to actin and actin cytoskeleton as well as terms related to secretory vesicles and secretory granules (Figure 5C).

**Figure 5.**
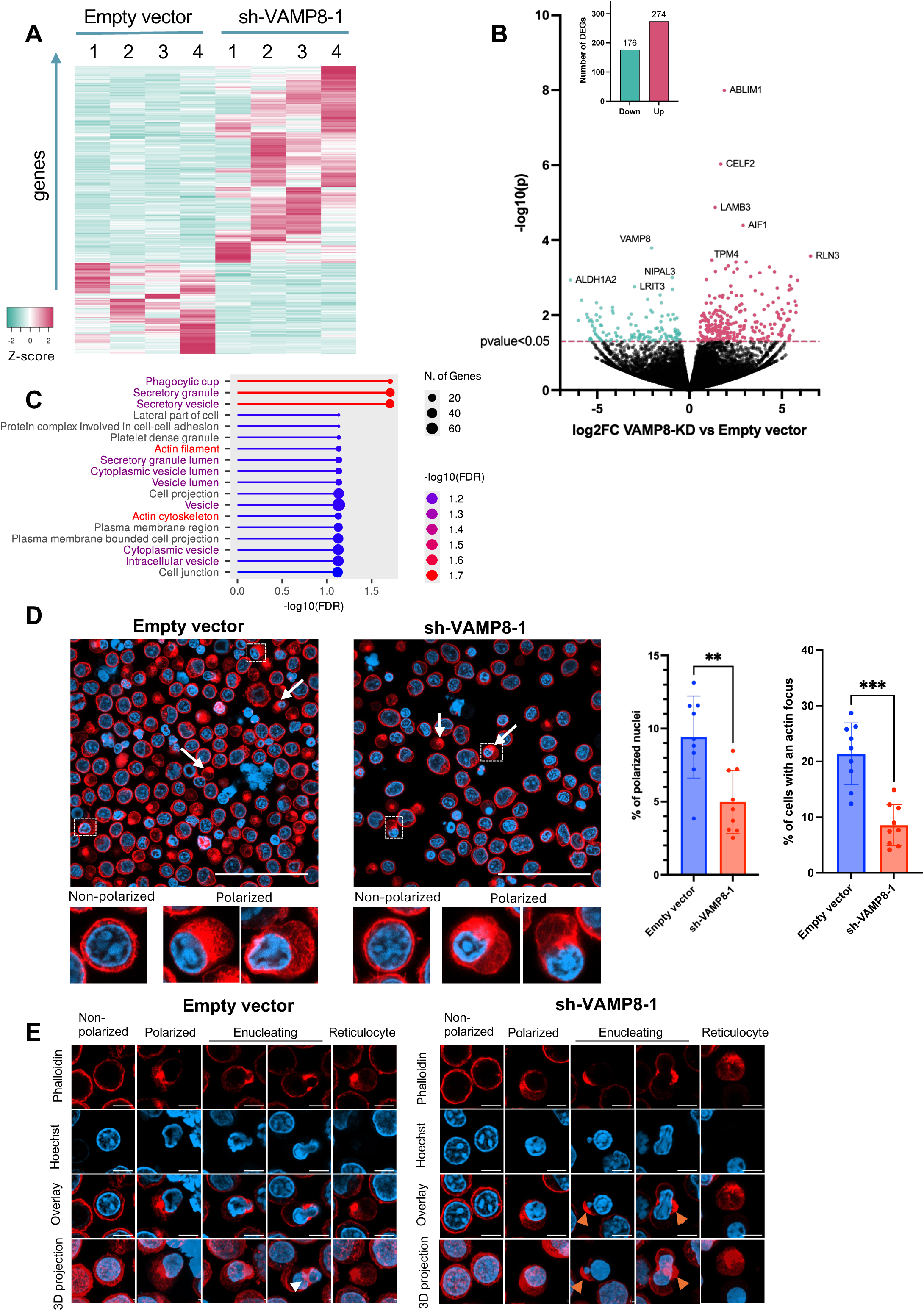
VAMP8-KD erythroblasts exhibit impaired nuclear polarization and disordered actin. (A) Heatmap depicting differentially expressed genes in WT (Empty vector) versus VAMP8-KD (sh-VAMP8-1) day 11 erythroblasts. The color scale indicates *z*-score of gene expression. (B) Volcano plot of differentially expressed genes in isogenic WT versus VAMP8-KD day 11 erythroblasts. *n*=4 biological replicates for each genetic background. (C) Pathway analysis for differentially expressed genes between WT and VAMP8-KD day 11 erythroblasts, using the Gene Ontology data base. (D) Immunofluorescence assays for localization of actin and cellular nuclei in isogenic WT or VAMP8-KD day 14 erythroblasts. F-actin was detected by phalloidin stain and nuclei were detected by Hoechst stain; scale bar, 50 μm. White arrows indicate actin foci. White boxes highlight cells with polarized or non-polarized nuclei, which are magnified 5X in the insets. Bar plots show quantification of nuclear polarization (left) and distinct F-actin foci (right). Data are from all cells across N=9 fields of view (>700 cells per condition). *P* -values were calculated using a Mann-Whitney test (** *P*-value <0.01; *** *P* -value<0.001). (E) Confocal slices of erythroblasts at various stages of enucleation stained with phalloidin and Hoechst. 3D projections derive from 39 confocal z-stacks for each picture. White arrows indicate a structure consistent with an actin ring in the control condition. Orange arrows point to aberrant actin structure. Scale bar, 5 µm.

Nuclear polarization is a key step during enucleation, and is proposed to depend on endosomal vesicular trafficking^44^ and actin reorganization^5^. To determine if VAMP8 is important for nuclear polarization, we performed immunofluorescence assays in in WT versus VAMP8-KD cells on day 15. We found that the VAMP8-KD cells had a significantly lower rate of nuclear polarization compared to the control cells (Figure 5D; supplemental Figure 5A).

While murine enucleation has been described as propelled by a contractile actin ring (CAR)^45^, studies in human cells have primarily highlighted a more diffuse F-actin focus at the site of the enucleosome ^5,6^. Using primary cRBCs derived ex-vivo from human bone marrow HSPCs, we observed an F-actin focus during nuclear polarization in both genetic backgrounds. Yet its frequency was significantly reduced in VAMP8-KD cells compared to WT, a difference that remained after nuclear extrusion (Figure 5D; supplemental Figure 5B-D). In the WT cells, we observed some cells with organized actin structures at the extrusion site, reminiscent of the contractile actin ring described in murine models (Figure 5E). This observation suggests a structure consistent with a CAR as part of the human enucleation process. In contrast, while we observed actin foci in VAMP8-KD cells, we did not detect any cells with organized ring-like structures. Instead, enucleating VAMP8-KD cells were often characterized by misshapen nuclei and disorganized actin aggregates (Figure 5E). Together, these results suggest that VAMP8 plays an important role in orchestrating the actin rearrangements required for enucleation, potentially including formation of the enucleosome.

## DISCUSSION

The phenomenon of erythroid enucleation is critical to the formation of healthy red blood cells, yet the mechanisms controlling this process have remained largely mysterious, in part because of the technical challenge of studying cells without nuclei. Here, we developed a forward genetic CRISPR-Cas9 screen in enucleated RBCs derived from primary HSPCs, leading to the discovery of novel regulators of enucleation. Key to our approach was the use of a modified CROPseq vector, where the sgRNA cassette is embedded into the 3’ LTR, enabling incorporation of the guide sequences into exported mRNA, which was retained in the cytoplasm after enucleation. Another important factor for successful pooled screening relates to the knockout efficiency. In our studies, we found that changing the scaffold to that of LRG2.1, which increases guide expression by disrupting the run of 4 thymidines found in the original CROPseq design^27,46,47^, substantially increased the knockout efficiency obtained with SLICE, where Cas9 is delivered by nucleofection, separately from lentivirus transduction. Further optimization of the kinetics of sgRNA and Cas9 delivery in these primary cells could maximize the probability and duration of their interactions and enhance the power of future screening efforts.

Notably, interpreting screens in dynamic, differentiating biological systems such as ex-vivo erythropoiesis requires careful consideration. In our screen, VAMP8-targeting sgRNAs were enriched in the enucleated cRBCs relative to unsorted cells, suggesting that VAMP8 may negatively regulate enucleation. However, characterization of the VAMP8-KD cRBCs revealed a striking failure of enucleation and accumulation of late-stage orthochromatic erythroblasts, suggesting a role for VAMP8 in promoting (rather than inhibiting) enucleation. We attribute this paradox to a kinetic bias, where the faster maturation of VAMP8-depleted cells leads to greater mRNA transcript decay after 17 days, artificially suppressing the representation of functional VAMP8 sgRNA sequences in the unsorted population. This highlights the importance of functional validation of any screening hits and suggests that the other highly ranked genes in our dataset may similarly lead to new insights into erythroid enucleation.

Our screen revealed a novel role for CLIC3 in erythroid differentiation, expanding on its previously studied role in cell proliferation in cancer cells^33,34,48,49^. We found that *CLIC3* depletion leads to cell cycle dysregulation and a delay in erythroblast maturation; this correlates with the activation of p53 and its downstream target *p21*. Elevated p53 and p21 are hallmarks of ineffective erythropoiesis and congenital anemias, which is characterized by apoptosis of early progenitors^38,39^. However, the CLIC3-depletion phenotype appears distinct for several reasons. We observed minimal cell loss, and erythroblasts, although delayed, were able to progress to the late orthochromatic stage before failing to enucleate. A potential explanation is provided by the reported interaction between CLIC3 and the N-acetyltransferase NAT10. In bladder cancer, elevated CLIC3 has been associated with reduced p21 levels, with a proposed mechanism that its interaction with NAT10 inhibits the enzyme’s ability to write N4-acetylcytidine (ac4c) modification of *p21* mRNA, leading to destabilizaton^34^. In the absence of CLIC3, ac4c modification of *p21* was shown to be enhanced, leading to increased mRNA stability. Investigating whether the cell cycle dysregulation observed in CLIC3-KD cRBCs involves an interaction with NAT10 and changes to *p21* mRNA modification has the potential to expand our understanding of the fundamental mechanisms controlling terminal erythropoiesis. Future studies aimed at determining if p21 and/or p53 inhibition can rescue the CLIC3-KD phenotype may also serve to clarify the underlying mechanism.

The SNARE protein VAMP8 was also identified in the screen as an important regulator of enucleation. Given VAMP8’s known role in autophagosome-lysosome fusion^42^ and the importance of organelle clearance for enucleation^3^, we first hypothesized that this block was due to a defect in organelle clearance. However, our finding that mitochondrial clearance progressed normally in the VAMP8-KD cRBCs suggest that VAMP8 is not required for this process, distinguishing its function from proteins like VDAC1^43^ and suggesting a non-canonical role for VAMP8 beyond autophagy.

Functionally, we showed that VAMP8-depleted cells have reduced efficiency of nuclear polarization, coupled with failure to form organized F-actin structures at the extrusion site. Our RNA-seq analysis also revealed an enrichment for terms related to secretory vesicles and granules. This suggests that VAMP8’s role here may not be degradative autophagy, but rather regulating trafficking of specific vesicles, such as those transporting critical polarity factors or membrane components to the nascent cleavage furrow. The failure of this targeted delivery could disrupt the establishment of polarity and, prevent the proper assembly of the actin machinery required for nuclear expulsion. Thus, VAMP8 may have a role at the intersection of vesicular trafficking and cytoskeletal organization; future studies will be important to dissect this complex and unexpected role during human enucleation.

The identification of CLIC3 and VAMP8 provides new insights into the molecular mechanisms regulating erythropoiesis and enucleation, highlighting a role for CLIC3 in cell cycle regulation and a role for VAMP8 in actin dynamics. Functional investigation of the other top candidates that arose from the screen, such as *CLU*, *CAND1*, *APP*, and *FARSA* has the potential to similarly advance our understanding of the complex process of erythroid enucleation. Moreover, forward genetic CRISPR screening in enucleated cells as introduced here may be applicable to a variety of biological processes and diseases that have been historically difficult to study, including reticulocyte maturation, platelet development, and infectious diseases such as malaria and babesiosis.

## Acknowledgments

The authors thank Jasmine Moshiri, Barbaro Baro, Melanie Espiritu, Ramesh V. Nair, the Stanford Shared FACS Facility, and the Stanford PAN Facility for technical assistance, and the Genetic Perturbation Platform at the Broad Institute for generating the CRISPR library and performing sequencing. The authors thank Lars Steinmetz for their CROPSeq library, and to the de-identified bone marrow donors for primary cells used in this study. The authors are grateful to John Boothroyd, Matthias Garten, Prasanna Jagannathan and members of their labs, members of the Egan Lab, and Alexander Marson, Eric Shifrut, Jake Freimer, Lars Steinmetz, and Daniel Schraivogel for helpful discussions.

This work was supported, in part, by NIH grants DP2HL13718601 (E.S.E.) and R01HL166249 (E.S.E). M.T. and R.C.G. were supported by The Katharine McCormick Advanced Postdoctoral Scholar Fellowship to Support Women in Academic Medicine and M.T. was also funded by a postdoctoral fellowship from the Stanford Maternal Child Health Research Institute. E.S.E. is a Biohub-San Francisco investigator and a Tashia and John Morgridge Endowed Faculty Scholar of the Stanford Maternal and Child Health Research Institute.

## Authorship

Contribution: M.T. and E.S.E. conceptualized the study; M.T., T.L., N.A.P., and R.C.G. performed experiments; all authors performed data analysis; J.G.D. and E.S.E. provided resources; M.T. and E.S.E. wrote the manuscript; all authors edited the manuscript; E.S.E. supervised the study and acquired funding.

## Correspondence

Elizabeth S. Egan, Stanford University School of Medicine, 240 Pasteur Drive, BMI 2400, Stanford CA 94305;

## Supplemental Figure Legends

**Figure S1. sgRNAs encoded in CROPseq lentivirus are incorporated into mRNA and are detected in the cytoplasm after erythroblast enucleation.** (A) PCR to detect LentiGuide-Puro-expressed sgRNAs in day 15 nucleated or enucleated cRBCs. (B) Schematic of sgRNA expression from the CROPseq lentiviral construct, the sgRNA cassette is embedded in the 3’ LTR, leading to its incorporation into a polyadenylated transcript that may be exported to the cytoplasm as well as generation of an RNA polymerase III sgRNA for genome editing. (C) PCR analysis to detect sgRNAs in day 15 nucleated and enucleated cRBCs transduced with CROPseq lentivirus expressing an sgRNA targeting GYPA. (D) Correlation plot of normalized sgRNA read counts between the day 8 population and the day 17 enucleated population for 39 unique sgRNAs.

**Figure S2. Forward genetic CRISPR-Cas9 screen in enucleated cRBCs identifies *CLIC3* and *VAMP8* as candidate determinants of erythroblast enucleation.** (A) Flow chart of the design of a custom sgRNA library targeting the erythrocyte proteome. The published erythroid proteomes from four studies were combined and filtered for proteins that were found in at least two studies. Essential genes and proteins required for erythropoiesis were excluded. The final library included 5007 sgRNAs targeting 681 genes (7 guides/ gene) and 240 non-targeting guides. (B) Volcano plots showing gene-level enrichment (median of LFC for all sgRNAs per gene) in enucleated vs. unsorted cells on day 17 for each individual replicate. Genes for which sgRNAs were significantly under-represented in the enucleated population (negative hits) are in blue, and genes with increased abundance (positive hits) are in pink. (C) Relative abundance of seven individual sgRNAs targeting CLIC3 in day 8, day 17 unsorted, and day 17 enucleated populations across three biological replicates. Individual sgRNAs are indicated by different colors. (D) Scatter plot showing the correlation in sgRNA abundance in the day 17 enucleated population vs unsorted population. The non-targeting sgRNAs are depicted in grey and the CLIC3 sgRNAs are in color (E) Scatter plot showing the correlation in sgRNA abundance in the day 17 Unsorted population vs the day 8 population. (F) Relative abundance of individual sgRNAs targeting CLIC3 in enucleated vs. unsorted cells. Each dot represents the mean ± SD for a single sgRNA of the three biological replicates. *P*-values were calculated using an unpaired t test (\**P*-value <0.05). (G) Scatter plot showing the correlation of sgRNA abundance in the enucleated population vs the unsorted population. (H) Relative abundance of individual sgRNAs targeting VAMP8 in enucleated vs. unsorted cells. Each dot represents the mean ± SD for a single sgRNA in three biological replicates.

**Figure S3. Depletion of CLIC3 in differentiating cRBCs leads to dysregulation of cell cycle genes, including those in the p53 transcriptional network.** (A) Volcano plot showing differentially expressed genes between WT and CLIC3-KD erythroblasts on day 11 shown in with differentially expressed genes within the p53 regulation network highlighted in teal. (B) Representation of the p53 transcriptional regulation network, from *Wiki Pathway* (WP4963). Genes found to be differentially expressed in CLIC3-KD compared to isogenic WT cells are highlighted in red. Differentially expressed genes were observed in several categories of the p53 transcriptional regulation, such as cell cycle arrest, apoptosis, and stem cell function. (C) Bar plot of p53 mRNA levels in day 11 CLIC3-KD erythroblasts (sh-CLIC3-1) and isogenic WT (Empty vector), as measured by qRT-PCR. Data are normalized to GAPDH and expressed relative to WT. Values represent the mean ± SD of *n*=2 biological replicates.

**Figure S4. VAMP8-KD cRBCs undergo mitophagy similar to WT during terminal erythroid differentiation.** (A) Gating strategy to measure mitochondrial load in specific erythroblast populations. Day 14 cRBCs were first gated for the presence of a nucleus, then based on their alpha4-integrin/Band3 profiles, and finally for Mitotracker staining to quantify mitochondria. (B) Representative experiment showing histograms of the mitochondrial loads in WT and VAMP8-KD erythroblasts at the Baso E, Poly E, or Ortho E stage of differentiation.

**Figure S5. Approach for quantification of nuclear polarization.** (A) Representative immunofluorescence images showing the strategy used to quantify nuclear polarization (Figure 5D). Nine fields of view were counted; yellow dots with “1” represent nucleated cells with a non-polarized nucleus and green dots with “2” represent nucleated cells with a polarized nucleus (defined as a nucleus positioned asymmetrically against the cell membrane). (B) Barplot showing the quantification of the percentage of cells with an actin focus among the cells with a polarized nucleus in isogenic WT (Empty vector) or VAMP8-KD (sh-VAMP8-1) as counted in N=9 pictures. *P*-values were calculated using an unpaired t-test (\**P*-value<0.05). (C) Barplot showing the quantification of the percentage of cells with an actin focus among the reticulocytes in isogenic WT (Empty vector) or VAMP8-KD (sh-VAMP8-1). *P*-values were calculated using an unpaired t-test (\**P*-value<0.05).

