## Supplemental Figures for "A pooled CRISPR screen reveals genes critical for erythroblast enucleation"

### Supplemental Figure 1

**A**

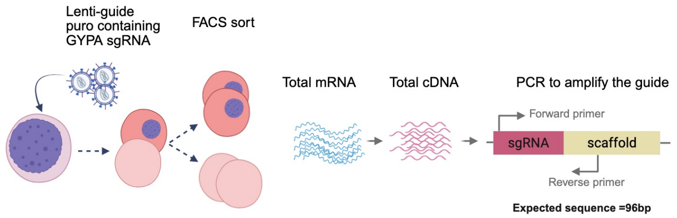


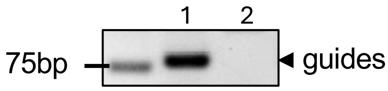


1: Nucleated cRBCs
2: Enucleated cRBCs

**B**

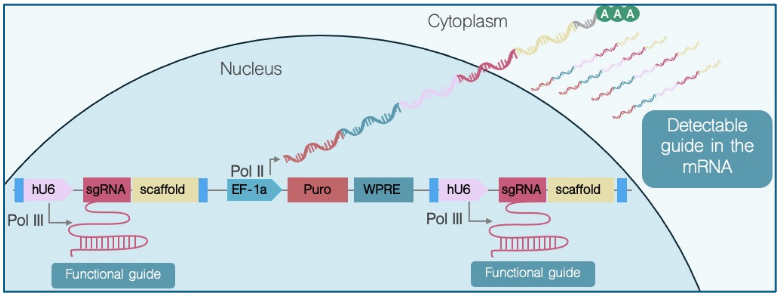


**C**

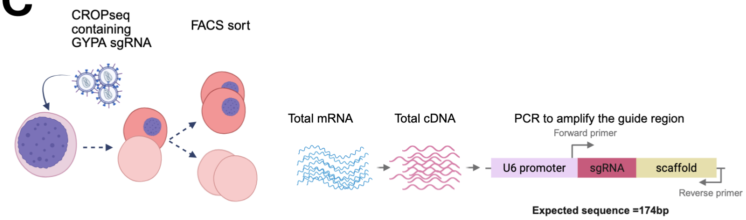


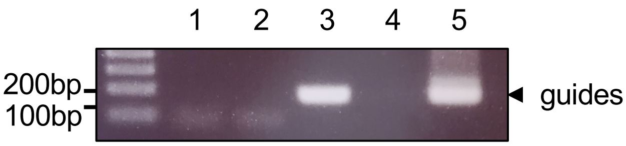


1: Enucleated, non transduced
2: Nucleated, non transduced
3: Enucleated + CROPseq
4: Empty well
5: Nucleated + CROPseq

**D**

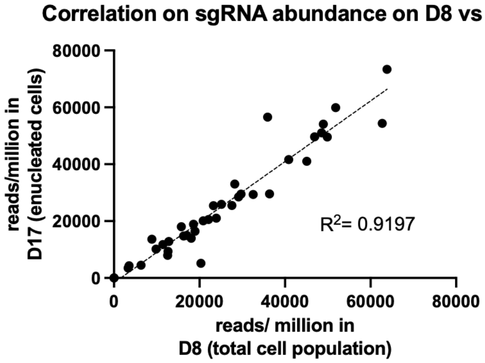

### Supplemental Figure 2

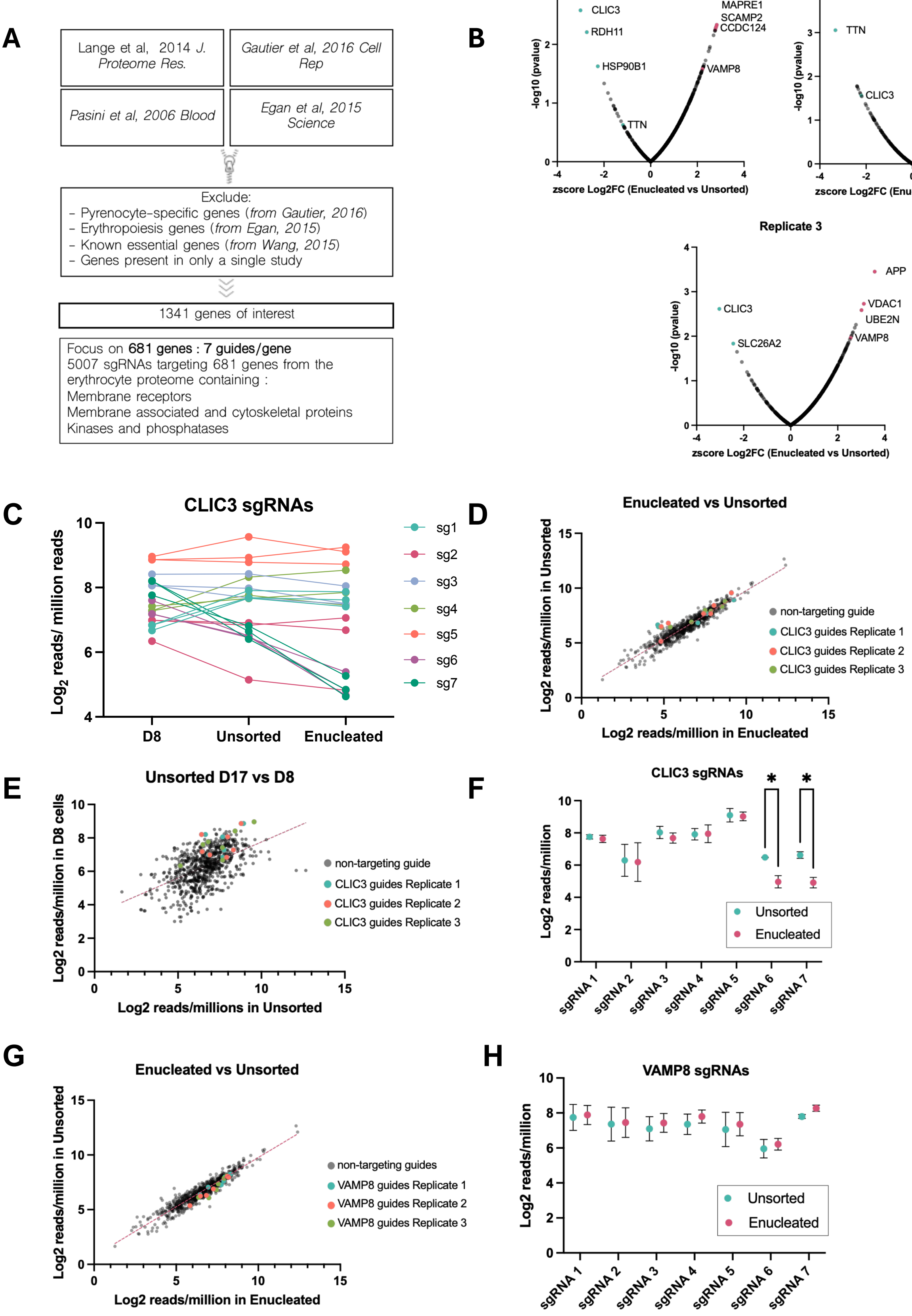

### Supplemental Figure 3

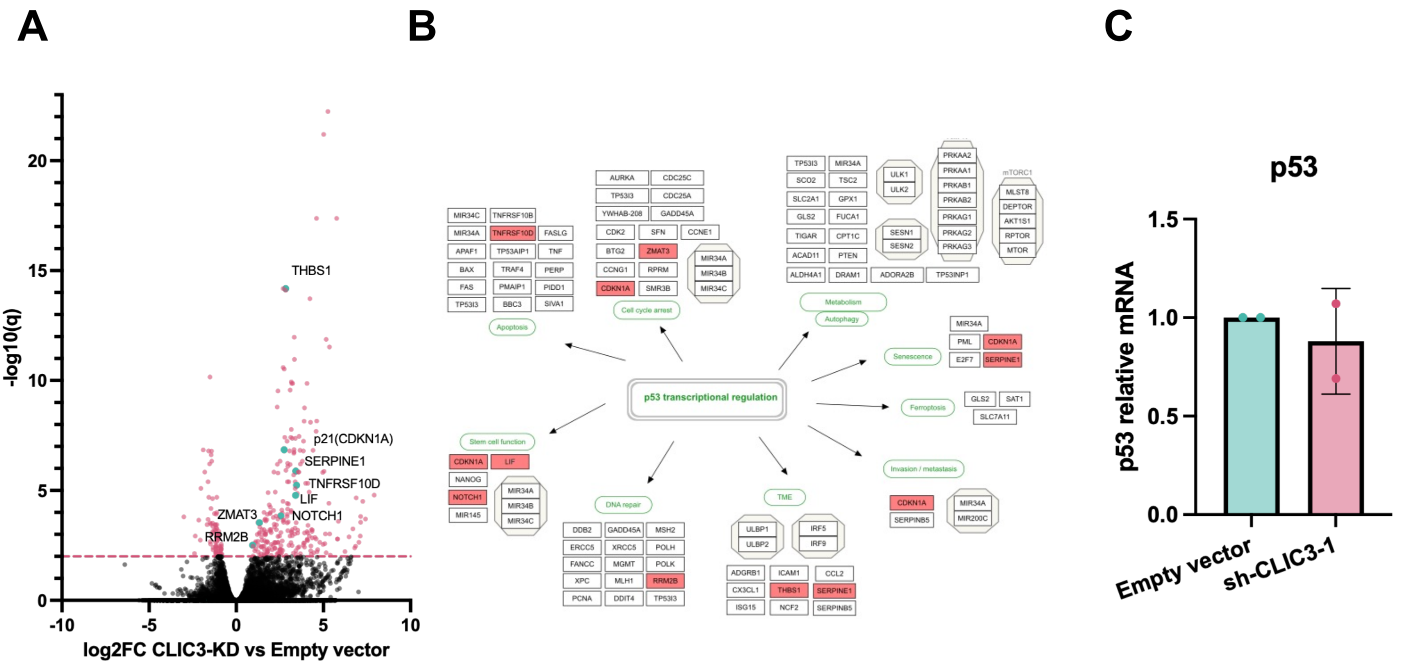

### Supplemental Figure 4

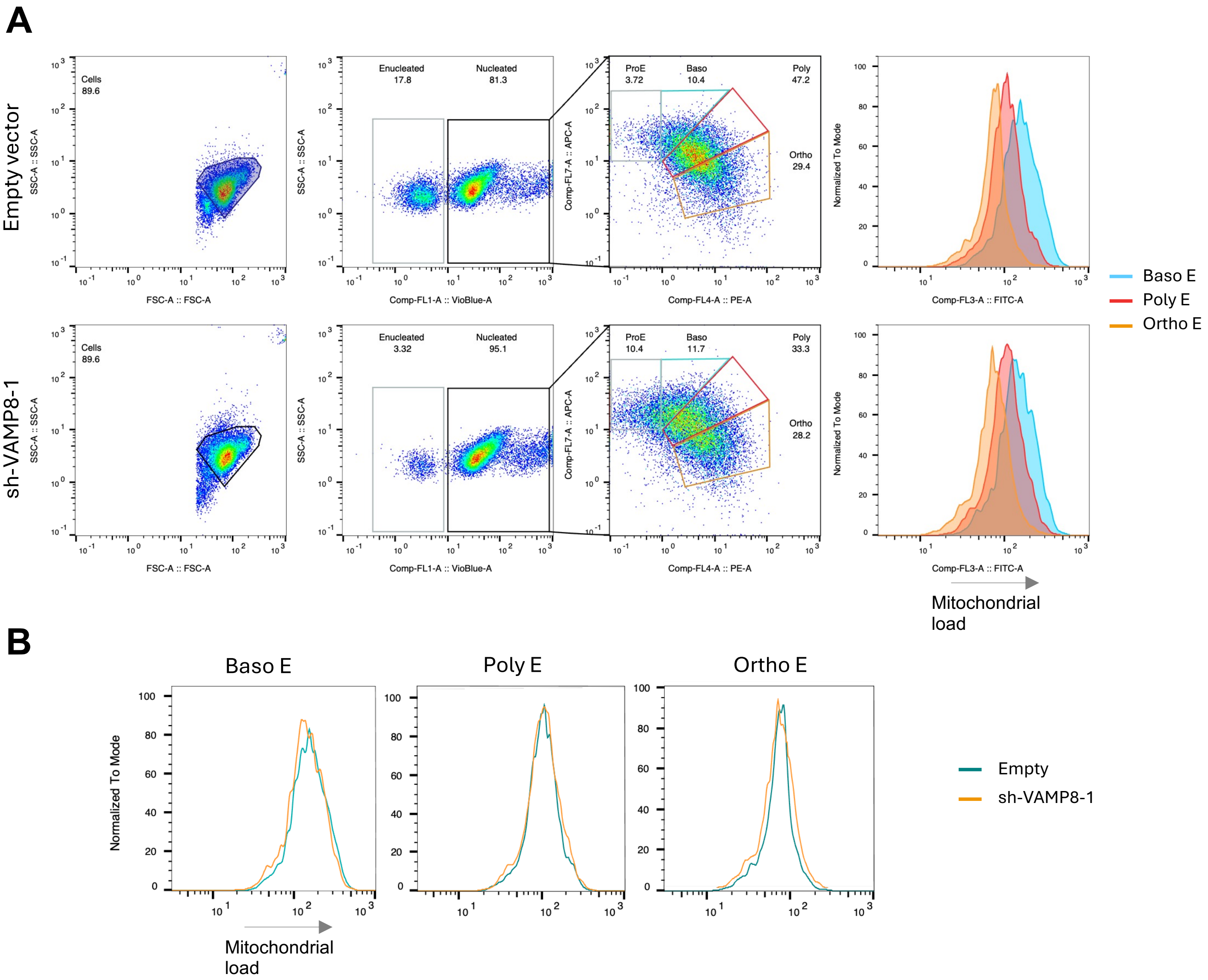

### Supplemental Figure 5

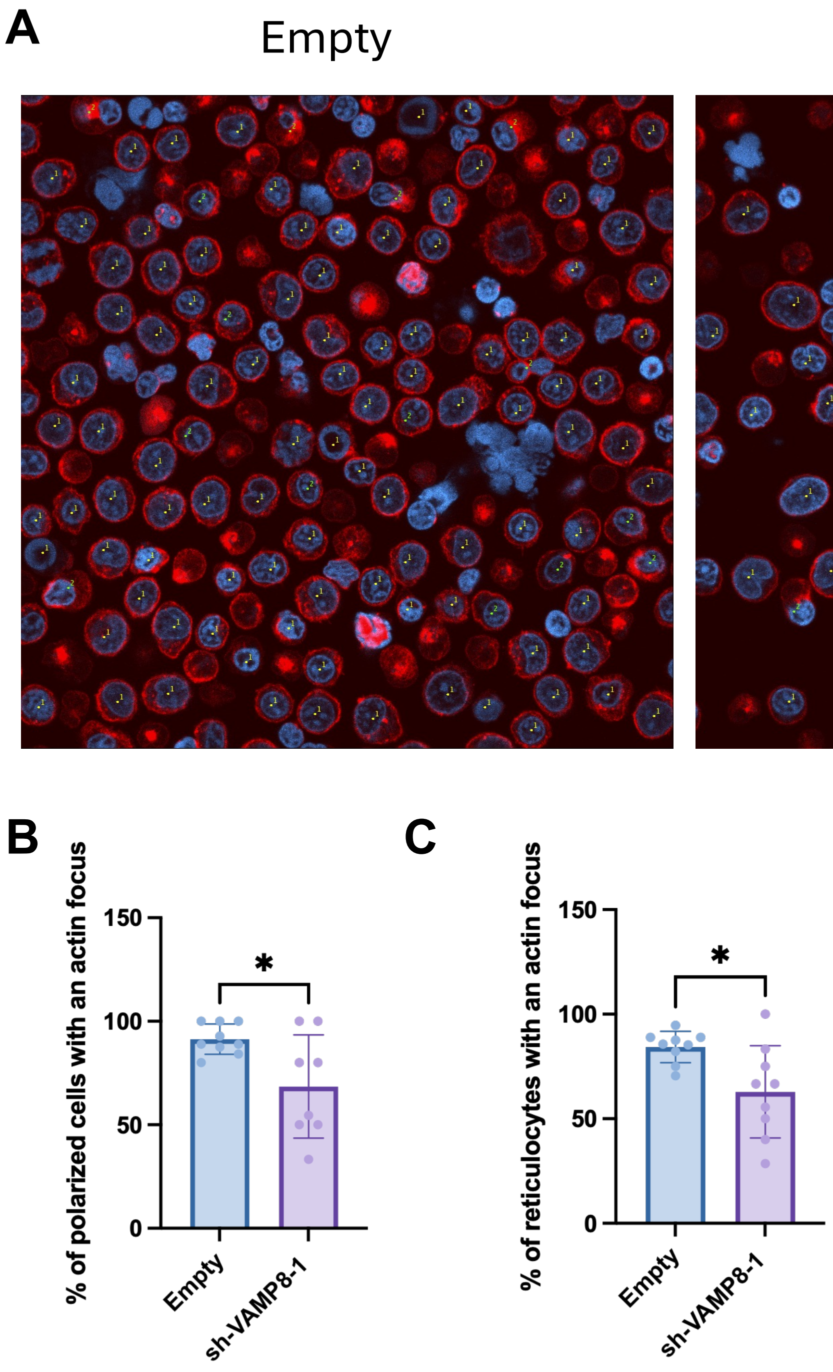
